# Fitness-Dependent Recombination Can Be Evolutionarily Advantageous in Diploids: a Deterministic Mutation–Selection–Balance Model

**DOI:** 10.1101/381228

**Authors:** Sviatoslav Rybnikov, Zeev Frenkel, Abraham B. Korol

**Affiliations:** Institute of Evolution, University of Haifa, Haifa 3498838, Israel; Department of Evolutionary and Environmental Biology, University of Haifa, Haifa 3498838, Israel; Department of Mathematics and Computational Science, Ariel University, Ariel 40700, Israel

**Keywords:** recombination, fitness dependence, diploids, purifying selection, recombination modifier

## Abstract

Recombination’s omnipresence in nature is one of the most intriguing problems in evolutionary biology. The question of why recombination exhibits certain general features is no less interesting than that of *why it exists at all*. One such feature is recombination’s fitness dependence (FD). The so far developed population-genetics models have focused on the evolution of FD recombination mainly in haploids, although the empirical evidence for this phenomenon comes mostly from diploids. Using numerical analysis of modifier models for infinite panmictic populations, we show here that FD recombination can be evolutionarily advantageous in diploids subjected to purifying selection. This advantage is associated with benefits from the differential rate of disruption of lower- *vs* higher-fitness genotypes, that can be manifested in systems with at least three selected loci. We also show that in systems with linked modifier, an additional contribution to the evolutionary advantage of FD recombination may come from fitness-dependence of the intensity of modifier linkage to the selected system, although the contribution of the last effect vanishes with tighter linkage within the selected system. We also show that in systems with three selected loci, FD recombination may give rise to negative crossover interference, which may be beneficial by itself. Yet, the role of such FD-induced crossover interference in the evolutionary advantage of FD recombination is minor. Remarkably, FD recombination was often favored in situations where any constant non-zero recombination was rejected, implying a relaxation of the rather strict constraints on major parameters (e.g., selection intensity and epistasis) required for the evolutionary advantage of non-zero recombination formulated by classical models.

## Introduction

Meiotic recombination is a process of reshuffling the parental genetic material, which takes place when a sexual organism produces its gametes. For about a century, recombination’s omnipresence in nature has been a most intriguing question, given the evolutionary ambiguity of this process (Weismann, 1889; Fisher, 1930; Maynard Smith, 1971; Bell, 1982). Indeed, on the one hand, the new allelic combinations generated in this process serve as raw material to meet selection demands. On the other hand, it can also break down the existing combinations, including even the most successful ones. However, the question of why recombination exhibits certain general features is no less interesting than the initial question of why recombination exists at all (Korol *et al.*, 1994; Lenormand *et al.*, 2016). One such feature is recombination’s sensitivity to external and/or internal conditions affecting the proportion of recombinants in the progeny. Harold Plough was the first to show that recombination rates (RRs) can rise when the organism is exposed to an ecological stressor (Plough, 1917). Empirical studies provide accumulating evidence for the ecological plasticity of recombination (for recent reviews, see Bomblies et al. 2015; Modliszewski and Copenhaver 2017; Stapley et al. 2017). Earlier, it was noticed that recombination’s ecological plasticity is genotype-specific (Elliott, 1955; Wilson, 1959; Nakamura, 1966), although the pattern of such specificity remained obscure. In 1986, Zhuchenko et al. (1986) demonstrated that stressor-induced changes in RRs can be modulated by genotype fitness in a negative-feedback manner, so that less stress-tolerant genotypes show more pronounced increases in RR. Moreover, RR may be sensitive to fitness even when no ecological stressors are imposed, i.e., when variation in fitness among individuals results from their differential genetic background (e.g., deleterious mutations) rather than from their differential stress tolerance (Tucić *et al.*, 1981). In general, one can think of ‘fitness-dependent’ (Zhuchenko *et al.*, 1986) or ‘fitness-associated’ (Agrawal *et al.*, 2005) recombination; herein, we use the former term, abbreviated as FD recombination. Empirical evidence for this phenomenon is still very limited, for both stressor-induced (Kilias *et al.*, 1979; Zhuchenko *et al.*, 1986; Korol *et al.*, 1994; Khlebova, 2010; Zhong and Priest, 2011; Jackson *et al.*, 2015; Hunter *et al.*, 2016; Aggarwal *et al.*, 2019) and mutation-induced (Tucić *et al.*, 1981; Tedman-Aucoin and Agrawal, 2012) changes in RR. Importantly, this evidence comes from diploids – mainly from fruit flies (Kilias *et al.*, 1979; Tucić *et al.*, 1981; Korol *et al.*, 1994; Zhong and Priest, 2011; Tedman-Aucoin and Agrawal, 2012; Jackson *et al.*, 2015; Hunter *et al.*, 2016), but also from plants, such as tomato (Zhuchenko *et al.*, 1986) and wheat (Khlebova, 2010).

An intriguing question is whether FD recombination can be considered as an evolvable phenotype. Analysis of natural populations infers that variation in RRs may indeed be adaptive (Ritz *et al.*, 2017). Theoretical models developed to date have clearly demonstrated the evolutionary advantage of FD recombination in haploids (Gessler and Xu 2000; Hadany and Beker 2003a,b; Agrawal et al. 2005; Wexler and Rokhlenko 2007). However, extending these results to diploids required additional specific assumptions, such as cis/trans effects (Agrawal *et al.*, 2005). In contrast, our recent study showed that FD recombination can also be evolutionarily advantageous in diploids under cyclical selection (Rybnikov *et al.*, 2017). The same result was obtained for another type of condition dependence, i.e., ecological plasticity of recombination, again for diploids exposed to cyclical selection (Zhuchenko *et al.*, 1985; Rybnikov *et al.*, 2017).

To address the discrepancy between the results obtained in the aforementioned diploid-selection models, i.e., between the positive result of Zhuchenko *et al*. (1985) and the negative result of Agrawal *et al*. (2005), it was suggested that more complex selection regimes, such as cyclical selection, favor FD recombination more than less complex ones, such as directional selection or mutation–selection balance (Agrawal *et al.*, 2005). Alternatively, as suggested in our recent paper (Rybnikov *et al.*, 2017), what may really affect the evolutionary advantage/disadvantage of FD recombination in the considered models is the presence or absence of variation in fitness among genotypes. Obviously, variation in fitness should concern only those genotypes in which recombination may affect the population structure in the next generation, i.e., genotypes heterozygous for at least two selected loci; in all other genotypes, RRs are ‘immaterial’, in terms of Otto and Barton (1997). Herein, as in our previous study (Rybnikov *et al.*, 2017), we refer to genotypes heterozygous for at least two selected loci as ‘recombination-responsive’. To further explore this assumption, here we examine the evolution of FD recombination in diploids under mutation–selection balance, which is a relatively simple selection regime compared with the cyclical selection. The wild-type genotype is assumed to have the highest fitness, while mutations at any locus are deleterious. If mutations at different loci affect fitness in a purely multiplicative way, recombination is known to be neutral. However, if the presence of a mutant allele(s) simultaneously at several loci decreases fitness more markedly (synergistic epistasis), then recombination can appear evolutionarily advantageous, since it facilitates purging mutant alleles from the population (Feldman *et al.*, 1980; Kondrashov, 1984; Charlesworth, 1990; Gabriel *et al.*, 1993; Barton, 1995; Lynch *et al.*, 1995; Otto and Barton, 1997; Otto and Feldman, 1997).

Under this scenario, we test whether FD recombination can be favored over constant RR. First, we examine the competition between FD recombination and the *optimal* constant RR. At that, we are interested, first and foremost, in situations with *zero* optimal constant RR (although those with intermediate optimal constant RR are also considered). Then, we generalize the question and examined the competition between FD recombination and its *equivalent* (in terms of the population mean RR) constant RR, regardless of whether this equivalent content RR is optimal or not. In all tests, we compare the competing recombination strategies using the modifier approach (Kimura, 1956; Nei, 1967), i.e., based on the dynamics of selectively neutral recombination-modifying alleles in an infinite panmictic population subject to diploid selection against deleterious mutations. It should be noted that comparisons between ‘asexual reproduction’ and ‘sexual reproduction without recombination’ on the one hand, and between ‘sexual reproduction without recombination’ and ‘sexual reproduction with recombination’ on the other, show that the effect of sex *per se* (i.e., of segregation) is much stronger than that of recombination (Charlesworth, 1990). Yet, the latter is no less important, bearing in mind that segregation is typically accompanied by recombination. Here, we study the evolution of recombination assuming obligate sexual reproduction with total panmixia, which seems realistic for many higher diploid eukaryotes.

## Models and Methods

### (a) Life cycle

We consider an infinite population of obligate sexual diploids with total panmixia. The life cycle includes random mating, selection at the diploid level, and meiosis resulting in gametes of the next generation. The generations do not overlap. Let *x*_*ij*_ be a diploid genotype made up of haplotypes *i* and *j*. Its frequency *p*^*s*^ after selection (as an adult) can be calculated based on its frequency *p* before selection (as a zygote) and its absolute fitness *W*:

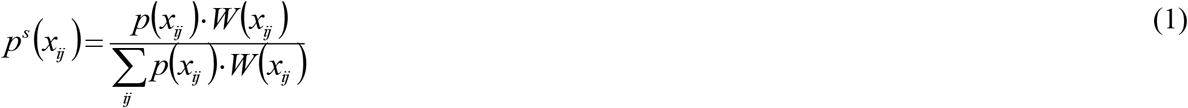

Then, let *g*_*k*_ be gamete of haplotype *k*. Its frequency *p* in the gamete pool can be calculated based on frequencies of adults and probabilities of recombination events:

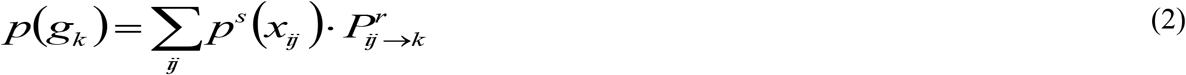

where 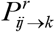 is the probability of obtaining gamete g_*k*_ from adult *x*_*ij*_ in meiosis, with the given rates of recombination and crossover interference.

Frequency *p*^*m*^ of a given gamete after mutation can be calculated based on probabilities of mutation events:

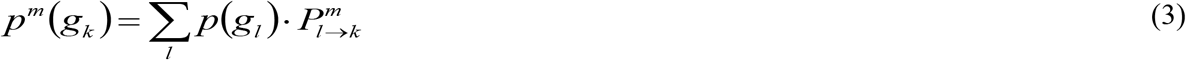

where 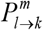 is the probability of obtaining gamete g_*k*_ from gamete g_*l*_ via mutations.

Finally, frequencies of zygotes in the next generation can be calculated based on frequencies of the corresponding gametes, given random mating:

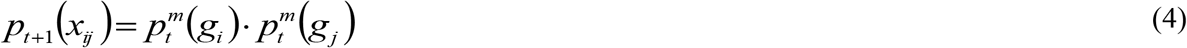

### (b) Genetic system and selection regime

Each genotype has either two (*A* and *B*) or three (*A*, *B,* and *C*) selected loci and a selectively neutral modifier locus (*M*) affecting RRs between the selected loci. The loci are arranged as *M*–*A*–*B*–*C*. Each selected locus is represented by two possible alleles: wild type (*A*, *B* or *C*) and mutant (*a*, *b* or *c*). The effect of the mutations on fitness is described by a standard multilocus model (Roze, 2009), as follows. The mutant alleles decrease fitness by *s* in the homozygous state and by *hs* in the heterozygous state (parameters *s* and *h* are referred to as ‘deleterious effect of mutation’ and ‘dominance of mutation’, respectively). For simplicity, both *s* and *h* are equal for all three selected loci. The interlocus interaction is multiplicative with epistasis (purely multiplicative selection is also considered as a specific case). If assumed, the epistasis is represented by three components: additive-by-additive (*e*_a×a_), additive-by-dominance (*e*_a×d_), and dominance-by dominance (*e*_d×d_), all of which are modeled as multiplicative terms. Thus, the fitness of the genotype bearing *N*_he_ heterozygous mutations and *N*_ho_ homozygous mutations is (Roze, 2009):

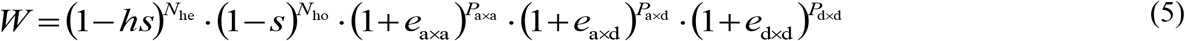

Here, the powers *P*_a×a_, *P*_a×d_ and *P*_d×d_ stand, respectively, for the number of additive-by-additive, additive-by-dominance and dominance-by-dominance epistatic interactions. These numbers can be obtained through *N*_he_ and *N*_ho_, as follows (Roze, 2009):

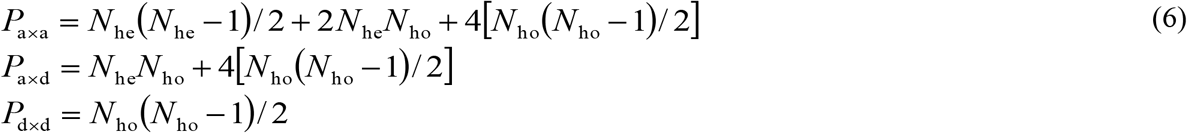

### (c) Recombination strategies

Modifier alleles define RRs within the selected system (*r*_*S*_). For simplicity, in the three-locus selected system, RRs between the adjacent selected loci are assumed to be equal (*r*_*AB*_ = *r*_*BC*_ = *r*_*S*_). The modifier locus is assumed to be either unlinked (*r*_*MA*_ = 0.5) or linked (*r*_*MA*_ = 0.05) to the selected system. The relations between modifier alleles are assumed to be purely co-dominant.

The modifier alleles confer various recombination strategies. We consider two types of strategies, implying that: (*i*) all genotypes of the selected system have the *same* RR (constant strategies, C-strategies), and (*ii*) different genotypes have *different* RRs, varying according to their fitness (FD-strategies). Under FD-strategy, RRs within the selected system (*r*_*S*_) depend negatively on genotype absolute fitness (*W*). Specifically, the genotype with the highest fitness (*W*_max_) has the lowest recombination rate (*r*_min_) and vice versa. For genotypes with intermediate fitness values, RRs are obtained by linear interpolation:

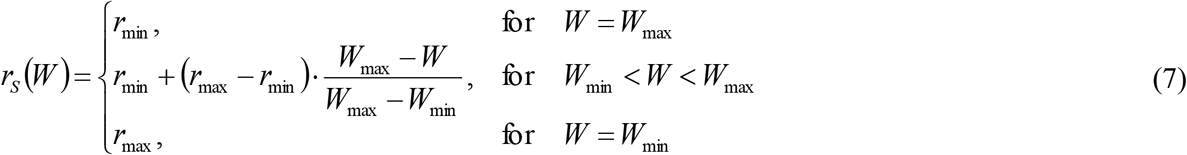

In our modeling, we assume that a precondition for the evolutionary advantage of FD recombination is variation in fitness among *recombination-responsive genotypes* (double and triple heterozygotes). In this respect, when estimating the lowest and highest fitness values (*W*_min_ and *W*_max_), we took into account only such genotypes. However, in models with two selected loci, there exists only one recombination-responsive genotype (double heterozygote), and such normalization would result in division by zero (equation 8). In this case, we estimated the lowest (*W*_min_) and highest (*W*_max_) fitness values among *all genotypes*.

In the majority of simulations, the magnitude of the plastic effect (Δ=*r*_max_−*r*_min_) was put equal to 0.05; we also examined higher magnitudes, up to 0.5.

### (d) Criteria for comparison of recombination strategies

First and foremost, we compared alternative recombination strategies in terms of *individual selection*, based on the dynamics of modifier alleles (Kimura, 1956; Nei, 1967). Specifically, strategy *S*_1_ was regarded as evolutionarily more advantageous than strategy *S*_2_ if the modifier allele for *S*_1_ succeeded in the two following tests. First, it had to invade populations in which the modifier allele for *S*_2_ prevailed. Second, it had to resist, when it itself prevailed, backward invasion by the modifier allele for *S*_2_. In both tests, ‘prevailing’ meant an allele frequency of 0.95. In both tests, modifier alleles were allowed to compete for 10,000 generations. Before the competition started, the selected system was allowed to evolve with a monomorphic modifier locus until it reached mutation–selection balance. The latter was diagnosed when allele frequencies at each selected locus changed by less than 10^−12^ per generation.

Aside from the dynamics of modifier alleles, we also compared recombination strategies in terms of the population mean fitness and population genetic variation. The population mean fitness was calculated as the fitness of all genotypes weighted by their frequencies. Population genetic variation (*v*) was calculated as the loci-averaged standard deviation of allele frequencies within the selected system:

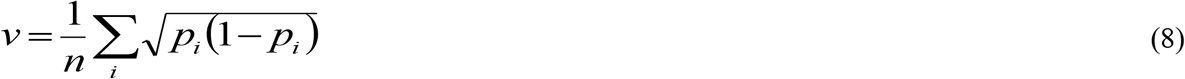

where *p*_*i*_ is allele frequency at the *i*-th selected locus, and *n* is the number of selected loci.

### (e) Design of the numerical experiments

We examined two selected systems: with two and three selected loci. For each of the two systems, we considered two situations: with unlinked (*r*_*MA*_=0.5) and linked (*r*_*MA*_=0.05) modifier, in order to address the potential effect of modifier linkage on the competition between alternative modifier alleles (for reviews, see Korol et al. 1994; Otto 2009).

For each of the four considered systems (with two and three selected loci, each with unlinked and linked modifier), we scanned ~600,000 combinations of five selection parameters: deleterious effect of mutations (*s*), the dominance of mutations (*h*), and three epistatic components (*e*_a×a_, *e*_a×d_, and *e*_d×d_). With respect to the deleterious effect, the mutations were scanned from almost neutral (*s*=0.01, as a proxy for *s*≈0) to lethal (*s*=1), with a step of 0.1. With respect to the dominance, the mutations were scanned from purely recessive (*h*=0) to purely dominant (*h*=1), with a step of 0.2. Due to the high difference between the results obtained for *h*=0 and *h*=0.2, we additionally examined the area of low dominance in more detail, with a step of 0.05. The epistatic components were scanned from –1 to 1, with a step of 0.1. Mutations were assumed to be unidirectional, with a rate of 10^−4^ per selected locus. No mutations were assumed at the modifier locus. In the first round of simulations, we assumed no crossover interference within the selected system (i.e. coefficient of coincidence *c*=1). Then, in order to test for the effect of crossover interference on the obtained results, we additionally examined two other situations: with full positive interference, implying no double-crossover events (*c*=0), and with considerable negative interference (with twice-higher frequency of double-crossover events, *c*=2). In these additional simulations, the additive-by-dominance epistasis was scanned with a step of 0.5, while the dominance-by-dominance epistasis was put equal to zero. The reason for treating the second (additive-by-dominance) and the third (dominance-by-dominance) epistatic components more roughly is that their effect, as we found in the first round of simulations, is, respectively, one and two orders of magnitude weaker compared to that of the first (additive-by-additive) epistatic component.

For each system and each combination of selection parameters, we compared FD recombination with two constant RRs: the optimal (for the given combination of selection parameters) constant RR and the equivalent (in terms of the population mean RR) constant RR. The optimal constant RR within the selected system 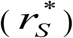 was estimated as follows. First, the minimal RR (*r*_*S*_ = 0) was compared with one step higher RR (*r*_*S*_ = δ*r*). If modifier allele for the latter invaded, it was compared with that for one more step higher RR (*r*_*S*_ = 2δ*r*), and so on. These pair-wise comparisons were conducted until modifier allele for a higher RR failed to invade. Once this happened, the previous RR was regarded as a lower estimate for the optimal constant 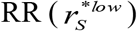. Second, we started with the maximal RR (*r*_*S*_ = 0.5) and moved downward (*r*_*S*_ = 0.5–δ*r*, *r*_*S*_ = 0.5–2δ*r*, etc.) until modifier allele for a lower RR failed to invade. Upon this, the previous RR was regarded as a higher estimate for the optimal constant 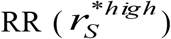. Then, we repeated the procedure between the obtained lower and higher estimates using the one order of magnitude smaller step, to obtain new, more accurate estimates. In total, we used three iterations, with steps equal to 0.01, 0.001 and 0.0001. The final lower and higher estimates differed by no more than 0.0001; the average between these two values was used as the optimal constant 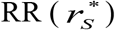.

The found optimal constant RR was then compared with FD recombination. In cases with *zero* optimal constant 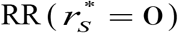, the latter was compared with FD-strategy implying RR variation from *r*_min_=0 to *r*_max_=Δ*r* (hereafter, we refer to such FD-strategy as ‘recombination-increasing FD-strategy’, or ‘+FD-strategy’). In cases with *intermediate* optimal constant 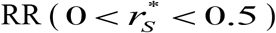, the latter was compared with three different FD-strategies implying RR variation: (*i*) from 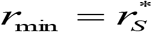 to 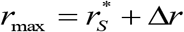 (‘recombination-increasing FD-strategy’, or ‘+FD-strategy’); (*ii*) from 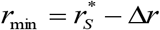 to 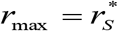 (‘recombination-decreasing FD-strategy’, or ‘–FD-strategy’); and (*iii*) from 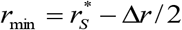 to 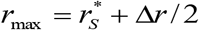 (‘fringe FD-strategy’, or ‘±FD-strategy’).

FD recombination was also compared with its equivalent (in terms of the population mean RR) constant RR. For a given interval, the population mean 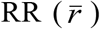 was calculated as the frequency-weighted RR in this interval:

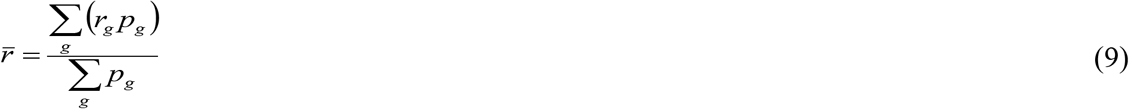

At that, only those genotypes were taken into account in which both loci flanking the studied interval were heterozygous. In this part of the experiment we considered, for each combination of the selection parameters, numerous FD-strategies, having the same fixed magnitude of the plastic effect (Δ*r* = 0.05) but differing in the range of RR (from *r*_min_ = 0 to *r*_max_ = 0.05, from *r*_min_ = 0.05 to *r*_max_ = 0.1, …, from *r*_min_ = 0.45 to *r*_max_ = 0.5).

The herein presented results are based on numerical simulations, which does not allow strict inferences about conditions that favor/disfavor FD recombination. Nevertheless, it is possible to discriminate parameter combinations leading to alternative outcomes (favor *vs* no favor) using numerical classification methods as a proxy, despite the fact that the considered model is purely deterministic and the described experimental design implies no stochasticity. Thus, to “quantify” the relative influence of the model parameters (the deleterious effect of mutations, their dominance, the epistatic components, the range of RRs under FD recombination, etc.) on the outcomes of the competition between FD recombination and constant RR, we employed the tools of logit analysis.

## Results

As expected, under *multiplicative* selection (*e*_a×a_ = *e*_a×d_ = *e*_d×d_ = 0), two arbitrary constant RRs always remained neutral to one another in terms of the modifier allele dynamics (i.e., the modifier alleles frequencies did not change regardless of their initial frequencies). Moreover, all constant RRs ensured the same equilibrium population mean fitness. Similarly, modifier alleles for FD recombination were neutral to those for the corresponding constant RR. This held for all examined systems: with two and three selected loci, and with the unlinked and linked modifier. Under *epistatic* selection, different constant RRs stopped being neutral one to another (in terms of both modifier-allele dynamics and population mean fitness), which allowed estimating the optimal constant RR. In the two following subsections, we present the results of competition between FD recombination and the corresponding (i.e., estimated for the same combination of the selection parameters) optimal constant RR, for cases with both zero and intermediate optimal constant RR. Then, in the third subsection, we pose a modified question and consider competition between FD recombination and its *equivalent* (in terms of the populations mean RR) constant RR, regardless of whether this equivalent constant RR is optimal or not within the class of constant RRs.

### (a) FD recombination *vs* zero optimal constant RR

As expected, in our simulations selection for/against non-zero RRs was strongly affected by the sign of the epistasis. Zero optimal constant RR was observed in a vast area of the parameter space: always under positive and often (but not necessarily) under negative additive-by-additive epistasis. The proportion of cases with zero optimal constant RR tended to decrease with a higher dominance of deleterious mutations, in accordance with the results reported by Roze (2009).

In the system with *two* selected loci, FD recombination was never favored over the corresponding zero optimal constant RR; this held for both systems with the unlinked and linked modifier locus. In contrast, in the system with *three* selected loci, FD recombination was non-rarely favored over the corresponding zero optimal constant RR. A necessary (but not sufficient) condition for the evolutionary advantage of FD recombination was negative epistasis. Other influential parameters, determining the outcome of the competition, were the deleterious effect of mutations and their dominance (Tables S1). In the system with *unlinked* modifier, FD recombination was favored under rather strong deleterious effects of mutations but intermediate negative additive-by-additive epistasis (so that the corresponding area of the parameter plane resembled a rightward-curved sickle). Low dominance of deleterious mutations tended to mitigate selection for FD recombination (Fig. 1A). In the system with *linked* modifier, zero optimal constant RR was observed in a smaller number of cases, typically under weak deleterious effects of mutations (unless the latter ones were purely recessive). However, almost in all such cases FD recombination was selected for, with the exception of those with weak negative additive-by-additive epistasis (Fig. 1B).

**Fig. 1.**
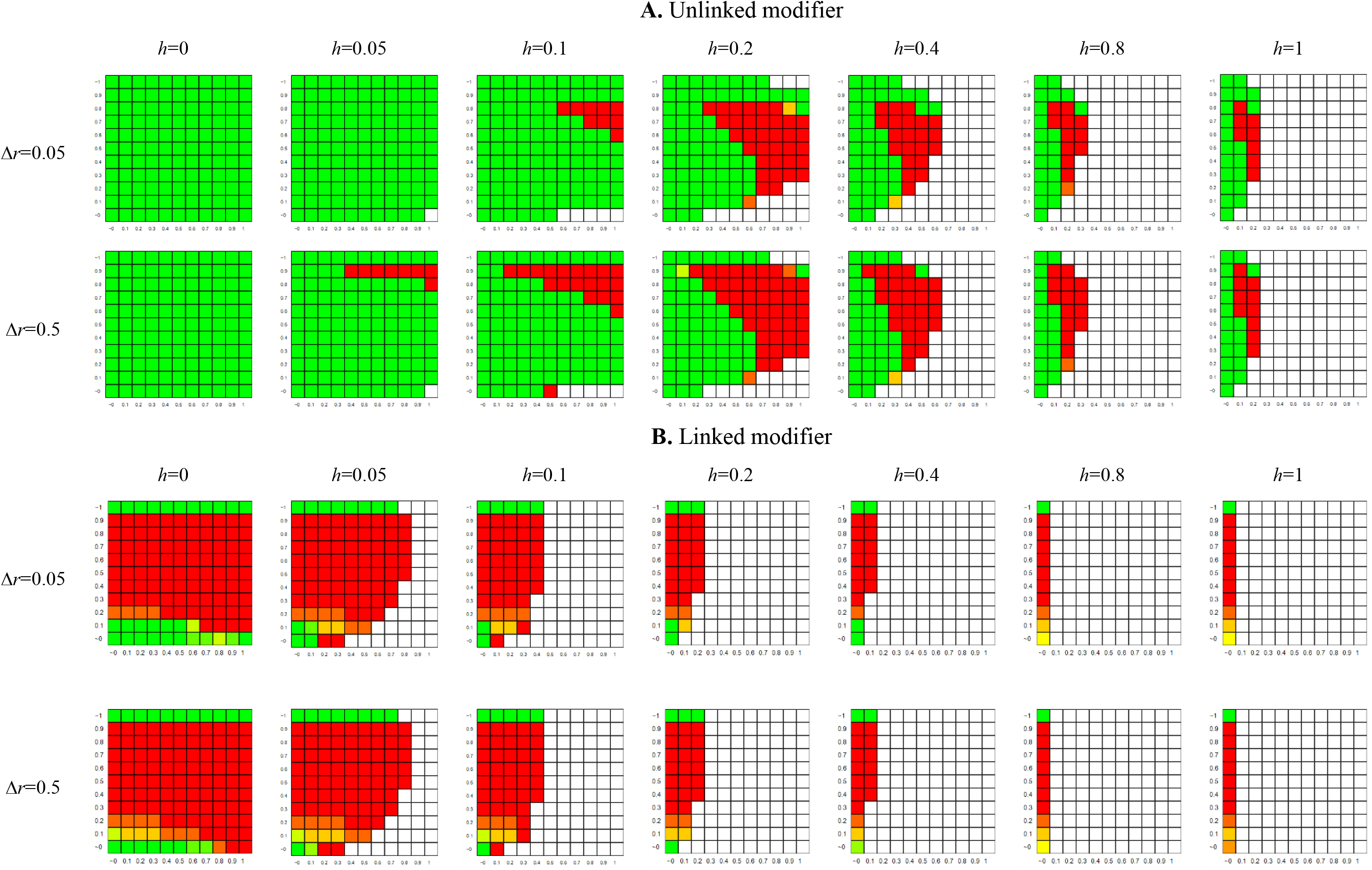
The evolutionary advantage of FD recombination over zero optimal constant RR: the effect of key selection parameters. The data stand for the system with three selected loci with no crossover interference within the selected system (however, the pattern appeared to be strongly robust to crossover interference – see below). Here and in Fig. 2–3, in each heat diagram, *x* and *y* axes stand, respectively, for the deleterious effect of mutations (*s*) and the absolute value of negative additive-by-additive epistasis (|*e*_a×a_|). The colors stand for the proportion of cases in which FD recombination was favored over zero optimal constant RR: from 0% (green) to 100% (red). No color means that no cases with zero optimal constant RR were observed under the given parameter combination.

In total, in the system with three selected loci, FD recombination was favored over zero optimal constant RR in ~23% and ~79% of cases with unlinked and linked modifiers, respectively. Perhaps, the higher proportion of cases favoring FD recombination in the system with linked modifier originates from a longer association of the modifier allele with the selected haplotypes; this longer association mitigates the constraints imposed on epistasis – as it happens when a population practices less sex compared with panmixia (Otto, 2009). With higher magnitudes of the plastic effect, the above-reported numbers increased, reaching ~30% and ~83% for unlinked and linked modifier, respectively. At that, higher magnitudes tended to relax the above-mentioned restriction on epistasis, making even strong negative additive-by-additive epistasis compatible with selection for FD recombination (Fig. 1).

Notably, although the overall number of cases in which FD recombination was favored grew with the magnitude of the plastic effect, in rare cases the high-magnitude +FD-strategy appeared less successful (in the competition against zero optimal constant RR) than the one with low-magnitude. This gave rise to the natural question of whether an *optimal* magnitude of the plastic effect exists. To address it, we let +FD-strategies with different magnitudes compete with one another and indeed found that in such cases a modifier allele for a certain intermediate magnitude displaced those for both smaller and larger magnitudes.

The above results were obtained for situations with no crossover interference within the selected system, which in principle should be considered as a special case rather than a rule. Therefore, we also examined two additional classes of situations: (a) with full positive interference, implying no double-crossover events (*c*=0); and (b) with negative interference, implying higher frequency of double-crossover events (we used *c*=2). The results appeared to be strongly robust. Specifically, the discrepancy in the proportion of cases in which FD recombination was favored over zero optimal constant RR compared to the situation with *c*=1 did not exceed 0.2%. Yet, crossover interference did affect quantitatively the competition between FD recombination and zero optimal constant RR: under lower values of *c*, the modifier allele for FD recombination invaded the population more easily.

We also compared the examined strategies (FD recombination and zero optimal constant RR) in terms of population-level characteristics. Under negative epistasis, FD strategy always increased the population mean fitness and always decreased the population genetic variation, regardless of whether FD recombination was favored or rejected. Yet, FD recombination led to much more pronounced changes in both characteristics (relative to those under zero optimal constant RR) when it was rejected rather than favored: ~10^−7^–10^−10^ against ~10^−9^–10^−13^ for the population mean fitness, and ~10^−3^–10^−6^ against ~10^−7^–10^−12^ for the population genetic variation. The changes were higher with the large magnitude of the plastic effect (by 1–2 orders compared to the changes obtained under the small magnitude) and with the linked modifier (by 2–3 orders compared to the changes obtained with the unlinked modifier). Apparently, whenever FD recombination was favored over zero optimal constant RR, it shifted upwards the population mean RR. Yet, in our simulations, this shift was very small: up to ~10^−5^–10^−3^ in the model with unlined modifier, and up to ~10^−4^–10^−2^ in the model with linked modifier. The reason is that the lower-fitness genotypes (i.e., those displaying higher RRs under FD recombination) remained very rare in the population subjected to purifying selection. As a consequence, selection for FD recombination also appeared to be very weak. We estimated that the invasion of the modifier allele for FD recombination can be stopped by burdening this allele with a multiplicative fitness-decreasing effect of ~10^−8^ –10^−10^ in the model with linked modifier and ~10^−10^ –10^−12^ in the model with unlinked modifier.

### (b) FD recombination *vs* intermediate optimal constant RR

Cases with intermediate optimal constant RR were found only under negative epistasis, as predicted by the theory (Feldman *et al.*, 1980; Kondrashov, 1984; Charlesworth, 1990; Gabriel *et al.*, 1993; Barton, 1995; Lynch *et al.*, 1995; Otto and Barton, 1997; Otto and Feldman, 1997). At that, the first (additive-by-additive) epistatic component was more influential than the second (additive-by-dominance) one, while the latter was more influential than the third (dominance-by-dominance) one. Specifically, selection for non-zero RRs occurred only under negative or at least zero additive-by-additive component. In the latter case, the next (additive-by-dominance) component had to be negative or at least zero. Finally, if both first (additive-by-additive and additive-by-dominance) components were zero, then the third (dominance-by-dominance) component had to be negative.

Whenever the optimal constant RR was intermediate, it was compared with three FD-strategies, namely with recombination-increasing, recombination-decreasing and ‘fringe’ ones (i.e., with RRs varying above, below and around the optimal constant RR, respectively). In the system with *two selected loci*, FD recombination was never favored. In contrast, in the system with *three selected loci*, the +FD-strategy appeared to be favored in a predominant proportion of cases (~76%), with the exception of marginal (either too low or too high) values of selection intensity and additive-by-additive epistasis. The evolutionary advantage of the ±FD-strategy was sporadic (~1%), and absent for -FD-strategy. Again, the above-presented results stand for the situation with no crossover interference (*c*=1). Simulations with full positive interference (c=0) and moderate negative interference (*c*=2) showed strong robustness of the parameter area in which FD recombination was favored over intermediated optimal constant RR, similar to the situations with zero optimal constant RR (Fig. 2). Noteworthy, negative interference considerably expanded the parameter area with intermediate optimal constant RR.

**Fig. 2.**
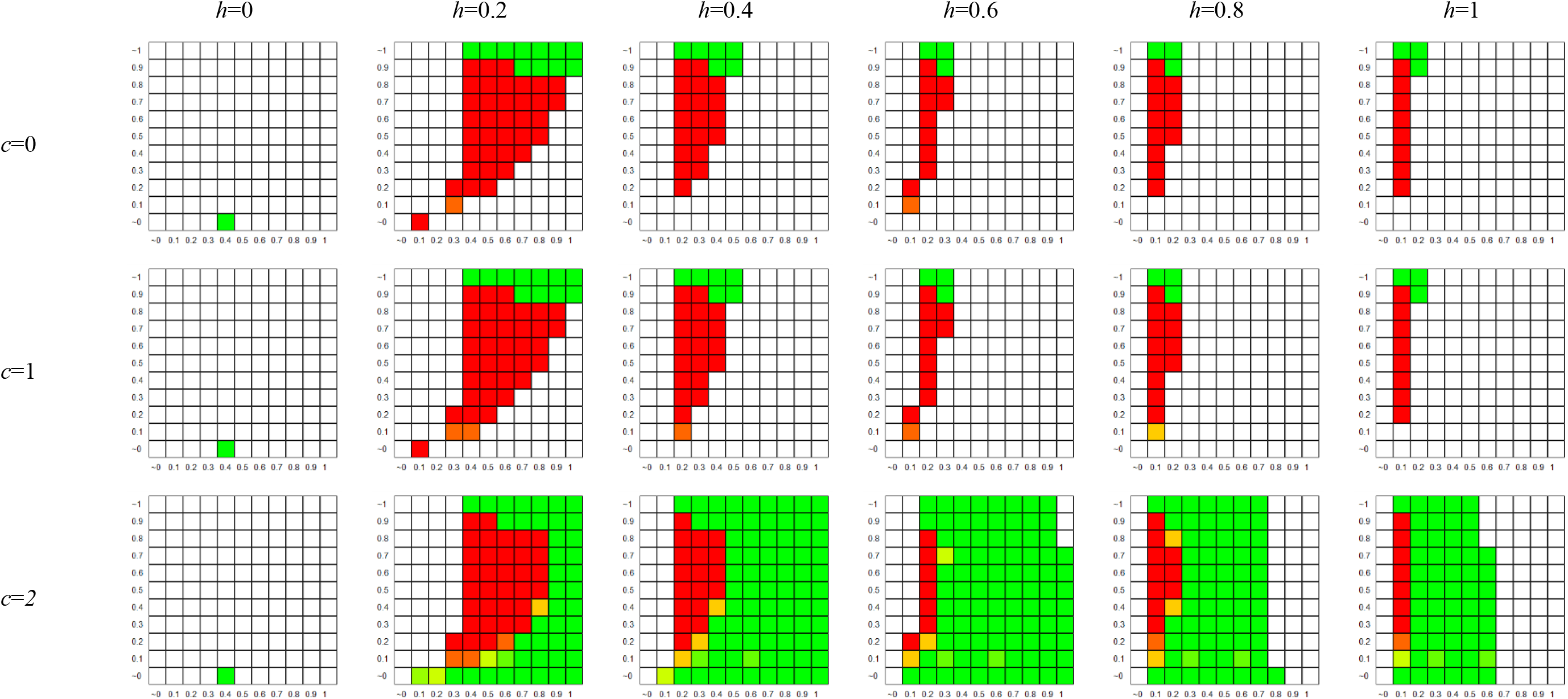
The evolutionary advantage of FD recombination over intermediate optimal constant RR: the effect of key selection parameters. The data stand for the system with three selected loci and linked modifier, +FD-strategy.

### (c) FD recombination *vs* equivalent constant RR

Apparently, examining competition between FD recombination and the *optimal* constant RR is attractive since it allows revealing situations in which FD recombination is favored over *all* constant recombination strategies (at least, unless the optimality is distorted by bi-stability or non-transitivity, which seem to be relatively rare phenomena). However, although populations evolve toward the optimal RR (Lobkovsky *et al.*, 2016), this evolution may be quite long, and a mutation in the modifier locus conferring FD recombination may appear much before the optimal constant RR is reached. With this respect, we found it reasonable to pose a question of whether FD recombination can be favored over its *equivalent* (in terms of the population mean RR) constant RR, regardless of whether this equivalent constant RR is optimal in the class of constant strategies or not. This question seems to be no less interesting than the traditional comparison between FD recombination and the optimal constant RR.

We found that in the system with *two* selected loci, FD recombination was always neutral in relation to its equivalent constant RR, but this neutrality disappears when we move from two to three selected loci. With *three* selected loci, FD recombination was quite often favored over its equivalent constant RR (Fig. 3). The most influential parameters, determining the outcome of the competition, were the deleterious effect of mutations, their dominance, and the additive-by-additive epistasis (similarly to what was observed in the competition between FD recombination and the optimal constant RR). RR within the selected system appeared to be influential only for the system with linked but not unlinked modifier (Table S2). In total, in the system with three selected loci, FD recombination was favored over its equivalent constant RR in ~59% and ~36% of cases with unlinked and linked modifiers, respectively. In the system with linked modifier, higher RR within the selected system tended to increase the proportion of cases in which FD recombination was favored: from ~27% for RR=0…0.05 up to ~44% for RR=0.45…0.5.

**Fig. 3.**
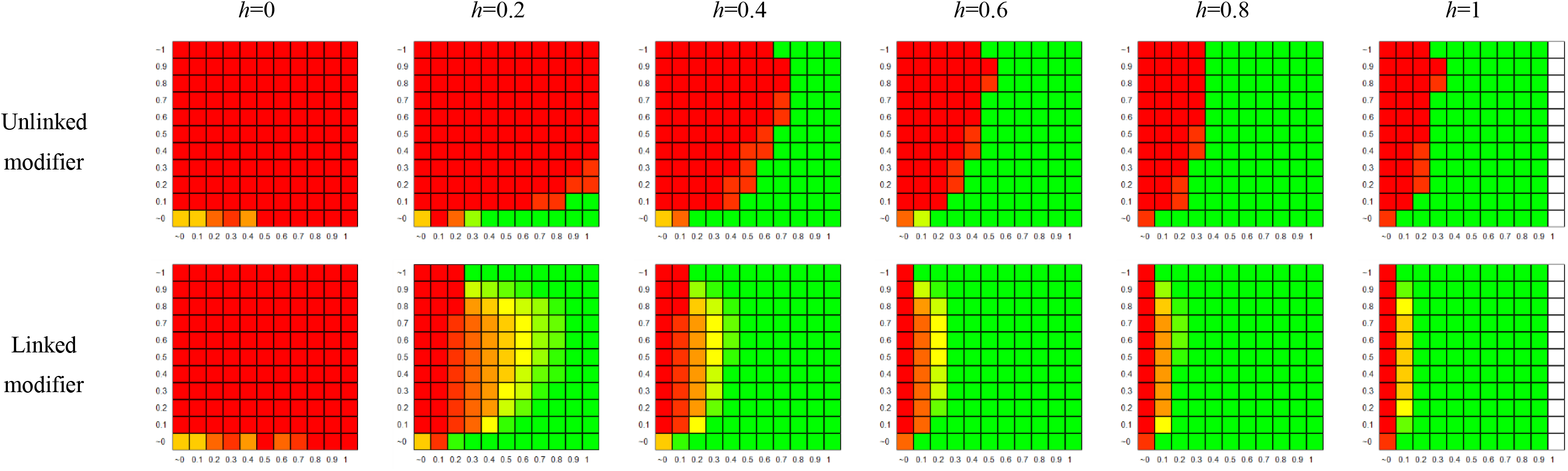
The evolutionary advantage of FD recombination over its equivalent constant RR: the effect of key selection parameters. The data stand for the system with three selected loci.

### (d) The role of FD-induced modulation of modifier linkage

In our simulations, we did not explicitly assume any effect of genotype fitness on the modifier linkage to the selected system. This means that RR between the modifier locus and the *adjacent* selected locus was always constant. However, RR between the modifier locus and *more distant* selected loci may become subject to an implicit variation. This happens under FD recombination in the system with three selected loci and linked modifier. Indeed, RR between the modifier locus and the second selected locus (i.e., *r_MB_*) depends on both *r*_*MA*_ and *r_AB_*; the variation in *r*_*AB*_ under FD recombination inevitably leads to some variation in *r_MB_*, even though *r*_*MA*_ is constant. This implicit variation in *r*_*MB*_ has an important consequence. The optimal constant RR within the selected system 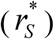 was estimated for a certain modifier linkage (*r*_*AM*_=0.05). However, under FD recombination, not all genotypes have this value of *r*_*AM*_. Thus, whenever we observed an evolutionary advantage of FD recombination over intermediate constant RR in the system with three selected loci and linked modifier, this advantage might have been associated, at least partly, with benefits from a ‘better’ mean RR (i.e., closer to the value which is optimal for the modifier linkage shifted due to *FD-induced* changes).

To address this issue, we additionally examined an FD-strategy implying the effect of genotype fitness only on *r*_*BC*_ but not *r*_*AB*_ (hereafter, it is referred to as ‘distant-interval’ FD recombination). For this strategy, we assumed twice higher magnitude of the plastic effect, in order to ensure a “more honest” comparability with the earlier considered ‘two-interval’ strategy. And indeed, the ‘distant-interval’ +FD-strategy appeared to be less advantageous over intermediate optimal constant RR than the ‘two-intervals’ +FD-strategy (~51% against ~76% of cases). At that, the difference between these two strategies tended to grow with the optimal constant RR in the distant interval (Fig. 4).

**Fig. 4.**
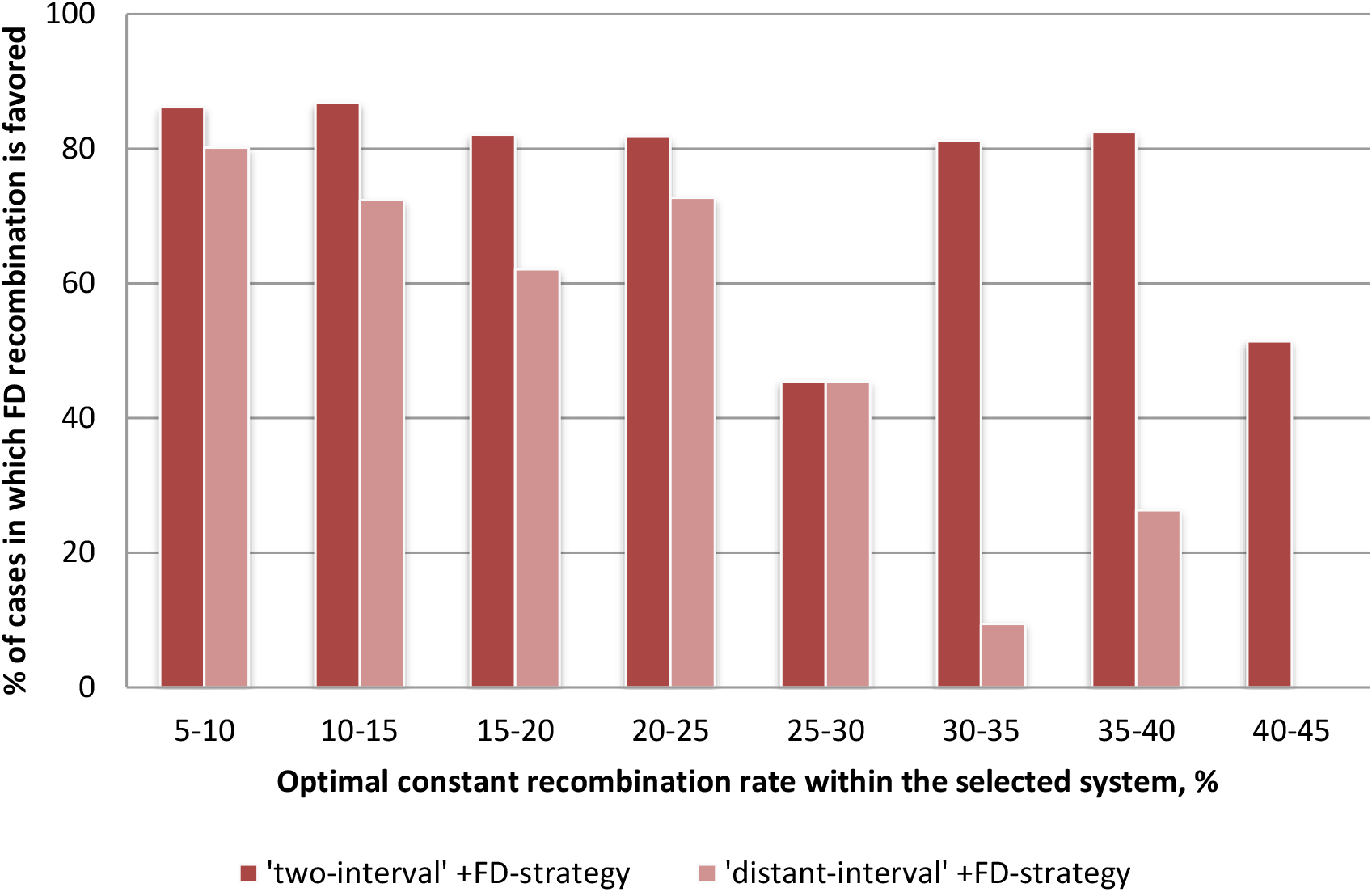
The role of FD-induced modulation of modifier linkage on the evolutionary advantage of FD recombination over intermediate optimal constant RR. The data stand for the system with three selected loci and linked modifier, +FD-strategy. The ‘distant-interval’ +FD-strategy can still be favored, although it is less advantageous than the ‘two-interval’ +FD-strategy. This suggests that FD-induced modulation of modifier linkage may play a certain role in systems with linked modifier, decreasing with linkage intensity between the selected loci.

### (e) The role of FD-induced crossover interference

When examining the competition between FD recombination and its equivalent constant RR, we found that under FD-strategy, the population mean RR in the derived interval deviates from that expected under the assumption of independence: 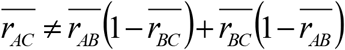. The value of such FD-induced interference-like effect can be estimated as follows: 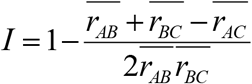. We found that FD recombination typically gives rise to a slightly negative interference-like values (*I* up to −0.02) (Fig. 5). Very rarely (in ~0.2% of cases), FD-induced interference was slightly positive (*I* up to 0.02); this happened under either very weak deleterious effect of mutations (*s*=0.01) or very weak additive-by-additive epistasis (*e*_a×a_=0.01) (Fig. 5).

**Fig. 5.**
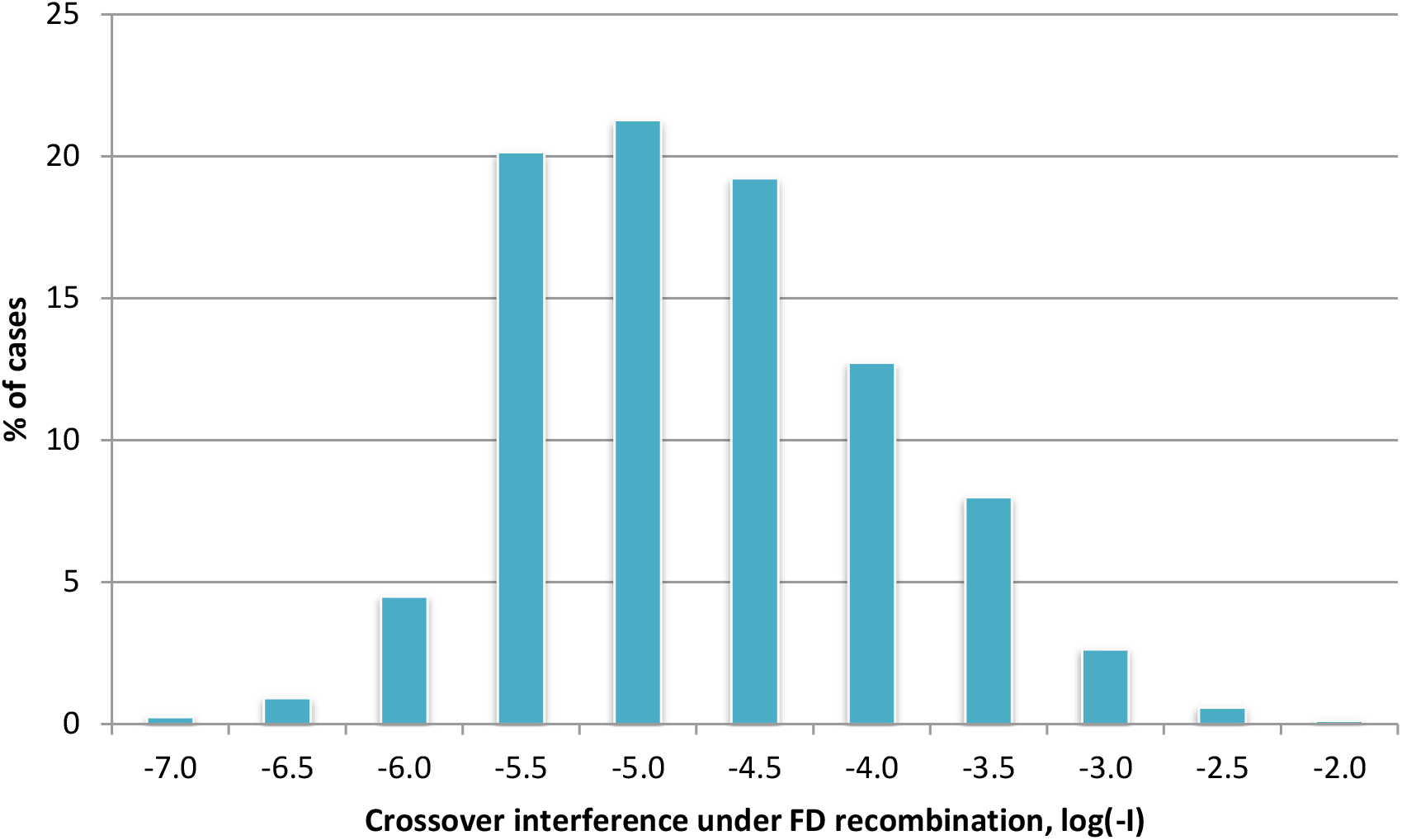
The distribution of cases by the value of FD-induced interference-like effect. Only cases with induced negative interference are shown. The numbers on *x*-axis stand for the left borders of the classes.

To test whether such FD-induced interference can be beneficial by itself, we compared two *constant* strategies: without crossover interference (i.e., equal to FD recombination in terms of *r*_*AB*_ and *r*_*BC*_ but not *r*_*AC*_) and with crossover interference (i.e., equal to FD recombination in terms of all three RRs: *r*_*AB*_, *r*_*BC*_, and *r*_*AC*_). We found that the modifier alleles conferring positive interference were never favored in this competition, while negative interference was advantageous quite often. However, the modifier-allele dynamics in the competition between the two constant strategies (with and without crossover interference) appeared to be very slow, ~1-2 orders of magnitude slower compared to the modifier-allele dynamics in the competition between FD recombination and its equivalent constant RR. Thus, in the system with three (and presumably, more) selected loci, FD recombination may give rise to an effect resembling crossover interference even though the simulation model did not assume interference explicitly. When being negative, such FD-induced interference may be advantageous by itself. However, even if such negative crossover interference is assumed for a constant strategy (what was actually done in the simulations described in subsection *c*), FD recombination is still favored. This suggests that the revealed evolutionary advantage of FD recombination goes beyond the advantage of negative interference-like effect induced by such strategy.

## Discussion

Several empirical studies have demonstrated that RRs are to a certain extent sensitive to genotype fitness, including situations in which the variation in fitness among genotypes originated from their different genetic backgrounds caused by deleterious mutations (Tucić *et al.*, 1981; Tedman-Aucoin and Agrawal, 2012). However, despite the progress in the theoretical explanation of the evolution of FD recombination in haploids (Gessler and Xu, 2000; Hadany and Beker, 2003a, 2003b; Agrawal *et al.*, 2005; Wexler and Rokhlenko, 2007), the so far suggested models encountered difficulties in extending the explanations to diploids.

We find it reasonable to distinguish between two forms of FD recombination, according to the ‘target’ genome region. The first one implies that genotype fitness affects only *RR between the modifier locus and the selected system* (so-called ‘selfish’ FD recombination). The evolution of the modifier allele for ‘selfish’ FD recombination is relatively easy to explain. Indeed, if a modifier allele is capable of somehow ‘evaluating’ its current genetic environment and tends to recombine from the low-fitness chromosome, it will inevitably increase its chances of surviving and spreading. Exploiting such benefits is now commonly referred to as the ‘abandon-ship’ mechanism (Agrawal *et al.*, 2005). Naturally, it requires the modifier locus to be linked to the selected system. However, the ‘abandon-ship’ mechanism was shown to be efficient in haploids (Hadany and Beker, 2003a; Agrawal *et al.*, 2005) but not diploids (Agrawal *et al.*, 2005). The intuitive reason is that in diploids, modifier alleles cannot ‘recognize’ which of the two homologous chromosomes causes lower fitness (unless the cis/trans effect, a rather specific phenomenon, is assumed). Thus, the modifier makes ‘right’ and ‘wrong’ decisions with equal probabilities and, therefore, with no effect on its allele frequencies in the next generation.

Importantly, in the current study, as well as in our previous one on the evolution of FD recombination under cyclical selection (Rybnikov *et al.*, 2017), we considered another form of FD recombination, implying that genotype fitness affects only *RRs within the selected system* (we call it ‘altruistic’ FD recombination, in opposition to ‘selfish’ FD recombination). We examined ‘selfish’ FD recombination only in an additional self-control experiment, aimed to test for the consistency of our models with those focusing on the ‘abandon-ship’ mechanism. In this additional experiment, the modifier allele for ‘selfish’ FD recombination was always neutral in relation to that for the optimal constant RR, which is consistent with the analytical solution of Agrawal *et al*. (2005). Moreover, two arbitrary modifier alleles were always neutral one to another if they conferred the same RRs within the selected system, regardless of their effect on RR between the modifier locus and the selected system. However, as already mentioned, our main simulations were conducted with ‘altruistic’ FD recombination. The key result is that ‘altruistic’ FD recombination can be favored in diploids, similarly to what was previously shown by Hadany and Beker (2003a) for haploids. However, the evolutionary advantage of this form of FD recombination is based on another mechanism. Specifically, if a modifier allele tends to disrupt lower-fitness selected genotypes more intensively than higher-fitness ones, it will increase the proportion of the latter in the population and thereby minimize, at least partially, the inherent costs of recombination.

In our simulations, ‘altruistic’ FD recombination was favored only in the system with three but not two selected loci. To our opinion, the reason is that the plasticity of RRs within the selected system implies not only benefits (differential rate of disruption of lower- *vs* higher-fitness genotypes) but also some costs. Indeed, the population mean RR established under FD recombination may depart from the optimal RR, which makes this new RR less favorable *by definition*. In such a situation, the fate of the modifier allele for FD recombination is determined by a tradeoff between the benefits and the costs of RR plasticity. While the costs emerge in both systems with two and three selected loci, the benefits require at least three selected loci. In the system with only two selected loci, the class of ‘recombination-responsive’ genotypes is represented by only one genotype - the double heterozygote (unless the cis-trans-effect is assumed). Obviously, this means no variation in fitness within this class, and hence *no fitness-associated variation* in RR. In fact, in the system with two selected loci, FD recombination degenerates into its equivalent constant RR, i.e., into the constant strategy with RR equal to that in the double heterozygote (this conclusion was confirmed by the explicit comparison between FD recombination and its equivalent constant RR, which showed their neutrality).

With respect to the above discussed tradeoff nature of the evolutionary advantage of FD recombination, we deduce that whenever FD recombination was favored in our simulations (in the system with three selected loci), the benefits of RR plasticity outbalanced its costs. Moreover, with the growing magnitude of the plastic effect, the benefits usually grew ‘faster’ than the costs (as reflected by the proportion of cases in which FD recombination was favored). In some cases, there existed an intermediate optimal magnitude, which suggests that the costs outbalanced starting from a certain threshold magnitude. This fact further argues that the evolutionary advantage of FD recombination is a kind of tradeoff. One more argument is the intriguing discrepancy between +FD-strategy and -FD-strategy in terms of their evolutionary advantage over the intermediate optimal constant RR: ~76% and <1%, respectively. Under purifying selection, the predominant part of the population is represented by the wild-type genotype (in our simulations, assuming a relatively modest mutation rate of 10^−4^ per locus, the mutant allele frequencies never exceeded 0.2%). Thus, the -FD-strategy (which decreases RR in higher-fitness genotypes) moves the population mean RR downwards much stronger than the +FD-strategy (which increases RR in lower-fitness genotypes) moves it upwards. Naturally, the strong departure from the optimal RR leaves almost no chances to -FD-strategy to be favored.

We argue that the key mechanism driving the evolution of FD recombination in diploids is the differential rate of disruption of lower- *vs* higher-fitness genotypes. However, FD recombination may give rise to some ‘by-product’ effects, which appeared to contribute to its evolutionary advantage. The first effect is FD-induced crossover interference. Our simulations confirmed that slight negative interference is often beneficial by itself, even in the class of constant RRs. Still, we demonstrated that FD recombination can be favored over the equivalent constant RR even if the latter is adjusted for crossover interference. An interesting question in this context is whether and how FD recombination can affect the evolution of crossover interference by itself, in line with the question earlier posed by Goldstein *et al*. (1993). The second effect is FD-induced modulation of the modifier linkage to the selected system. Contrary to FD-induced crossover interference, which occurs in any selected system with at least three loci, FD-induced modulation of the modifier linkage requires more specific conditions: (a) the selected system must consist of at least three linked loci, (b) the modifier locus must be linked to the selected system, and (c) at least three of the selected loci are located from one side of the modifier. Still, we demonstrated that in such situations FD recombination can be favored even if the effect of FD-induced modulation of the modifier linkage is excluded. Besides, FD recombination is often favored in the class of situations with no modifier linkage to the selected system.

The evolutionary advantage of FD recombination in diploids was observed in several population-genetics models: mutation-selection balance (the herein presented results), cyclical selection (Rybnikov *et al.*, 2017), and Red Queen dynamics (Rybnikov *et al.*, 2018), which argues for universality of the underlying mechanism. Remarkably, in all three mentioned population-genetics models FD recombination was shown to be favored under certain parameter combinations even if *any constant* non-zero RR was rejected. This indicates that assuming FD recombination enables selection for non-zero RRs under much milder constraints on key parameters (such as population size, selection intensity, epistasis, modifier linkage, etc.) compared to those revealed in a spectrum of classical models with constant RRs (Barton, 1995; Otto and Barton, 1997, 2001; Otto and Feldman, 1997). Thus, the amazing ‘recombination-supporting’ potential of FD recombination, first demonstrated by Gessler and Xu (2000) for haploids, can be extended also to diploids. Notably, the same pattern was revealed for another variation-affecting characteristic, the rate of sex, which is also long known to exhibit fitness dependence (for a recent review, see Ram and Hadany 2016). Specifically, FD sex was shown to be favored over asexual reproduction even if *any constant* non-zero rate of sex was rejected (Mostowy and Engelstädter, 2012).

As already mentioned, the ‘abandon-ship’ mechanism was shown to be efficient only in haploids but not diploids. At the same time, the differential rate of disruption of lower- *vs* higher-fitness genotypes can be evolutionarily advantageous in both haploids and diploids. We speculate that FD recombination first appeared in haploids. There, it could have been ‘invented’ by some ‘selfish’ genes, which spread by exploiting the ‘abandon-ship’ benefits (Gessler and Xu, 2000; Otto, 2009). Later on, such genes probably expanded the ability to affect RR in an FD mode to other genome regions. Once this happened, FD recombination stopped being entirely dependent on the ‘abandon-ship’ benefits, and could also evolve in diploids, where the ‘abandon-ship’ mechanism does not work alone. We consider such extension as a transformation of “*effect”* into “*function”* (Maynard Smith, 1982) in the course of evolution of recombination, fitting well the *relay-race* principle (Ratner, 1990).

We are aware of the limitations of simulation modeling and believe that the analytical treatment of the herein presented models will allow clarifying the underlying mechanisms. Still, we think that the obtained results, showing that the evolution of FD recombination in diploids is not a dead-end problem, warrant theoretical generalization and broad experimental studies.

## Acknowledgments

We thank Camille Vainstein for English editing of the manuscript. We are grateful to Aneil Agrawal and anonymous reviewers for their helpful critical comments on the first version of the manuscript. The work was supported by the Israel Science Foundation (grant number 1844/17), the Graduate Studies Authority of the University of Haifa, and the Ministry of Aliyah and Integration of Israel.

---

**Table S1:** The relative influence of model parameters on the evolutionary advantage of FD recombination over zero optimal constant RR (the system with three selected loci)

| Variable | Unlinked modifier |  | Linked modifier |  |
| --- | --- | --- | --- | --- |
|  | B | Wald | B | Wald |
| Deleterious effect of mutations ($s$) | 5.36 | 2536.66 | 0.85 | 67.49 |
| Dominance of mutations ($h$) | 5.34 | 2739.63 | 0.55 | 24.94 |
| Additive-by-additive epistasis ($e_{a \times a}$) | -1.22 | 268.01 | 0.19 | 4.63 |
| Additive-by-dominance epistasis ($e_{a \times d}$) | -0.02 | 0.88 | -0.10 | 7.30 |
| Crossover interference ($c$) | -0.05 | 4.15 | -2E-03 | 0.00 |
| Magnitude of the plastic effect ($\Delta r$) | 0.92 | 70.79 | 0.63 | 18.12 |
| Constant | -6.00 | 3148.97 | 1.03 | 135.29 |
| Total number of cases | 17205 |  | 8811 |  |
| Correctly predicted cases, % | 76.5 |  | 81.2 |  |

**Table S2:** The relative influence of model parameters on the evolutionary advantage of FD recombination over its equivalent constant RR (the system with three selected loci)

| Variable | Unlinked modifier |  | Linked modifier |  |
| --- | --- | --- | --- | --- |
|  | B | Wald | B | Wald |
| Deleterious effect of mutations ($s$) | -11.00 | 6228.56 | -6.74 | 6575.85 |
| Dominance of mutations ($h$) | -10.81 | 6484.09 | -7.22 | 7536.89 |
| Additive-by-additive epistasis ($e_{a \times a}$) | -3.94 | 2546.45 | -0.38 | 48.60 |
| Additive-by-dominance epistasis ($e_{a \times d}$) | -0.09 | 10.64 | -0.07 | 8.77 |
| RR within the selected system ($r_S$) | -0.40 | 6.42 | 3.42 | 613.82 |
| FD-induced crossover interference ($c$) | -3E-04 | 81.44 | -3E-04 | 49.53 |
| Constant | 10.23 | 5762.62 | 4.57 | 4076.40 |
| Total number of cases | 35736 |  | 35736 |  |
| Correctly predicted cases, % | 90.9 |  | 85.7 |  |

